# Functional maturation and experience-dependent plasticity in adult-born olfactory bulb dopaminergic neurons

**DOI:** 10.1101/2024.11.28.625840

**Authors:** Candida Tufo, Menghon Cheah, Marcela Lipovsek, Lorcan P Browne, Matthew S Grubb

## Abstract

Continued integration of new neurons persists in only a few areas of the adult mouse brain. In the olfactory bulb (OB), immature adult-born neurons respond differently to olfactory stimuli compared to their more mature counterparts, and have heightened levels of activity-dependent plasticity. These distinct functional features are thought to bestow unique properties onto existing circuitry. OB interneurons, including those generated through adult neurogenesis, consist of a set of highly distinct subtypes. However, we do not currently know the different cell-type-specific mechanisms underlying their functional development and plastic potential. Here, we specifically characterised electrophysiological maturation and experience-dependent plasticity in a single, defined subtype of adult-born OB neuron: dopaminergic cells. We selectively live-labelled both adult-born and ‘resident’ dopaminergic cells, and targeted them for whole-cell patch-clamp recordings in acute mouse OB slices. Surprisingly, we found that from the time – at ∼1 month of cell age – that live adult-born dopaminergic neurons could first be reliably identified, they already possessed almost fully mature intrinsic firing properties. We saw significant maturation only in increased spontaneous activity and decreased medium afterhyperpolarisation amplitude. Nor were adult-born dopaminergic cells especially plastic. In response to brief sensory deprivation via unilateral naris occlusion we observed no maturation-specific plastic alterations in intrinsic properties, although we did see deprivation-associated increases in spike speed and amplitude across all adult-born and resident neurons. Our results not only show that adult-born OB dopaminergic cells rapidly functionally resemble their pre-existing counterparts, but also underscore the importance of subtype identity when describing neuronal maturation and plasticity.

## Introduction

Adult neurogenesis continually produces new neurons that functionally integrate into existing circuitry in two major mammalian brain regions: the dentate gyrus of the hippocampus, and the olfactory bulb (OB; Lledo *et al*., 2006). In both regions, newly generated adult-born cells are different from their pre-existing neighbours by virtue of their transient immaturity. Before they become fully integrated into adult hippocampal or bulbar circuits, young newborn neurons possess distinct morphological and functional features, and have specialised response properties to sensory stimuli and behavioural tasks (Carleton *et al*., 2003; Espósito *et al*., 2005; Lledo *et al*., 2006; Whitman & Greer, 2007a; Grubb *et al*., 2008; Bardy *et al*., 2010; Livneh *et al*., 2014; Danielson *et al*., 2016; Wallace *et al*., 2017). Immature adult-born neurons are also more plastic than their mature counterparts, with an elevated capacity both for rapid synaptic plasticity and for longer-term alterations produced by chronic perturbations in ongoing activity levels (Schmidt-Hieber *et al*., 2004; Ge *et al*., 2007; Tashiro *et al*., 2007; Kelsch *et al*., 2009; Nissant *et al*., 2009; Gu *et al*., 2012; Livneh *et al*., 2014; Alvarez *et al*., 2016). These unique properties of immature, plastic newborn neurons are thought to underlie the specific contributions of adult neurogenesis to specific forms of learning and memory (Nakashiba *et al*., 2012; Grelat *et al*., 2018; Li *et al*., 2018; Bragado Alonso *et al*., 2019).

In the OB, the constitutive integration and maturation of adult-born neurons is characterised by significant cell-type diversity. Adult neurogenesis produces two major forms of GABAergic OB interneuron – granule cells and periglomerular cells – which not only populate different OB laminae and possess distinct molecular, morphological and functional identities, but also have their own distinctive maturational trajectories (Tufo *et al*., 2022). Moreover, within each of these major OB cell classes exists an impressive level of subtype heterogeneity, driven by the preferential production of different neuronal types from specialised spatial locations within the germinal region of the ventricular-subventricular zone (Merkle *et al*., 2007; Lledo *et al*., 2008; Fiorelli *et al*., 2015; Takahashi *et al*., 2016; Malvaut *et al*., 2017; Hardy *et al*., 2018; Mizrak *et al*., 2019; Cebrian-Silla *et al*., 2021). This diversity can be regulated by internal states such as food intake or pregnancy (Paul *et al*., 2017; Chaker *et al*., 2023), such that the subtype composition of adult-generated neurons is continually fine-tuned in response to ongoing organismal changes.

Diversity is particularly prevalent among adult-born OB periglomerular cells. These neurons assume non-overlapping molecular identities characterised by the mutually exclusive expression of the calcium-binding proteins calretinin or calbindin, or the dopamine-synthesising enzyme tyrosine hydroxylase (TH; Kohwi *et al*., 2007; Kosaka & Kosaka, 2007; Panzanelli *et al*., 2007; Parrish-Aungst *et al*., 2007; Whitman & Greer, 2007b). Postnatal production of these distinct periglomerular subtypes is known to be tightly regulated by an array of molecular factors (Hack *et al*., 2005; Kohwi *et al*., 2007; Brill *et al*., 2008; Cave *et al*., 2010; Ihrie *et al*., 2011; de Chevigny *et al*., 2012; Agoston *et al*., 2014; Bonzano *et al*., 2016; Tiveron *et al*., 2017; Coré *et al*., 2020; Remesal *et al*., 2020). The survival of adult-born periglomerular subclasses is also differentially regulated by alterations in sensory experience, with the dopaminergic TH-positive subpopulation especially responsive to long-term conditions of olfactory enrichment or deprivation (Bastien-Dionne *et al*., 2010; Sawada *et al*., 2011; Bonzano *et al*., 2014; Angelova *et al*., 2023).

However, despite this striking diversity, and despite the importance of immature properties in determining the role of adult-generated cells in existing circuits, the functional maturation and plastic potential of different adult-born OB periglomerular subtypes remain entirely unexplored. Non-specific targeting of newborn periglomerular cells has shown that, as an overall population, these neurons quickly develop basic functional properties which then mature steadily over an extended period lasting several months. In particular, synaptic inputs, spiking capabilities and sensory responses to odorant stimuli are all present in 1-2-week-old newborn periglomerular cells that have only just reached the glomerular layer. However, synaptic properties, spontaneous spiking, dendritic morphology, and odour selectivity do not fully match those of neighbouring ‘resident’ glomerular layer interneurons until around 6-9 weeks of age (Belluzzi *et al*., 2003; Mizrahi, 2007; Grubb *et al*., 2008; Livneh *et al*., 2009; Livneh & Mizrahi, 2011; Livneh *et al*., 2014; Kovalchuk *et al*., 2015; Liang *et al*., 2016; Maslyukov *et al*., 2018; Su *et al*., 2023). The overall population of adult-born periglomerular cells is also highly plastic over this extended timeframe, displaying alterations in dendritic morphology, synapse stability and odour response properties after prolonged manipulations of sensory input or intrinsic activity levels (Livneh *et al*., 2009, 2014; Livneh & Mizrahi, 2011; Li *et al*., 2023).

Our current knowledge of adult-born periglomerular cell maturation and plasticity, however, completely fails to account for the significant heterogeneity of this cell type. Might there be distinct maturational processes and plastic properties in individual adult-born periglomerular subclasses which may have been masked by treating these cells as a single population? The closest information we currently have on this question comes from studies using the location and intensity of TH-driven GFP expression as a proxy for the maturation status of adult-generated dopaminergic neurons. This approach has shown that weakly TH-GFP-positive cells in deeper layers of the OB – presumed to be very young dopaminergic-fated neurons migrating towards the glomerular layer – have immature electrophysiological properties and distinct gene expression patterns compared to strongly TH-GFP-expressing resident glomerular layer neurons (Pignatelli *et al*., 2009; Casciano *et al*., 2023). However, the potential for extended maturation and plasticity once adult-born OB dopaminergic cells are established in glomerular layer circuits remains unstudied.

Here we use a conditional expression approach to specifically live-label adult-born OB dopaminergic neurons, enabling us to target them, and their resident dopaminergic neighbours, for electrophysiological recordings at defined periods in their maturation. Surprisingly, we find that as soon as these adult-generated OB dopaminergic cells can be reliably identified when they are about 1 month of age, their intrinsic functional properties are already almost entirely mature. At this timepoint we also find no evidence for elevated levels of experience-dependent plasticity, although brief olfactory deprivation did induce significant alterations in spike strength in both adult-born and resident OB dopaminergic neurons alike.

## Methods

### Animals

All experiments were performed on adult mice of either sex, housed under a 12 h light-dark cycle in an environmentally controlled room with free access to food and water. DAT-tdTomato mice were generated by crossing DAT^IRESCre^ mice (B6.SJL-Slc6a3^tm1.1(cre)Bkmn/J^, Jax stock 006660) with a floxed-tdTomato reporter line (Gt(ROSA)26Sor^tm9(CAG-tdTomato)Hze^, Jax stock 007909). Wild-type C57BL/6N mice (Charles River) were used to back-cross each generation of transgenic mice, and for quantitative PCR experiments. All experiments were performed at King’s College London under the auspices of UK Home Office personal and project licences held by the authors.

### Stereotaxic Adeno-Associated Virus (AAV) injections

For live-labelling adult-born dopaminergic neurons, 2-3 month-old DAT-tdTomato mice were injected in the rostral migratory stream (RMS) with AAV8-CAG-FLEx-GFP virus (2 x 10^12^ gc/ml; UNC Vector Core). Prior to surgery for RMS virus injections mice were anaesthetised with an intraperitoneal injection of ketamine (Vetalar; 50 mg/Kg) and medetomidine (Domitor; 1 mg/Kg). The paw-pinch reflex and breathing were checked for sufficient depth of anaesthesia before mounting the mice in a stereotaxic frame with digital readout (World Precision Instruments, WPI). Carprofen (Rimadyl; 5 mg/Kg) was injected subcutaneously to relieve the mice from post-surgical pain, and eye lubricant gel was applied prior to the surgery. A skin incision was made above the skull to expose the surgical area and clippers were used to keep the surgical area exposed. One set of experiments used the following coordinates from Bregma: +3.3 mm AP; +0.82 mm ML (Grubb *et al*., 2008). However, these co-ordinates sometimes resulted in off-target labelling of clusters of GFP-positive cells in the caudal OB, likely due to a rostral ‘leak’ of AAV permitting direct infection and expression in resident, non-adult-born OB dopaminergic neurons. Such ‘leak’ label was readily identified upon visual inspection of acute slices, and any animals with this pattern of GFP label were omitted from our study. We subsequently minimised these cases by using more caudal injection coordinates: +2.55 mm AP; +0.82 mm ML (Hardy *et al*., 2018). At these coordinates small craniotomies were drilled over the RMS of the right hemisphere. After drilling the craniotomies, the same coordinates were taken again from Bregma to inject the virus, using a 33-gauge blunt custom-made needle (Hamilton) attached to a 10 µl syringe (RN701, Hamilton). In order to create a ‘pocket’ for the virus in the RMS and maximise absorption the needle was first lowered to -2.95 mm DV and immediately afterwards the virus was injected at -2.90 mm DV with a volume of 150 nl at 50 nl/minute through a microinjection pump with SMARTtouch controller (WPI). Three minutes after the injection was completed the needle was slowly removed from the brain. These three minutes were intended to minimise leak of virus from the pipette as it returned through more dorsal brain regions.

The surgical area was sutured and cleaned with an antiseptic skin cleanser, Hibiscrub. The mice were then given subcutaneous injections of saline to recover from blood loss and atipamezole (Antisedan; 5 mg/ml) to reverse the anaesthetic effect of the medetomidine. The mice were allowed to recover in a 32 °C chamber, then housed in either individual cages or together with siblings which underwent the same procedure.

### BrdU administration

For comprehensive labelling of newborn cells, 5-Bromo-2’-deoxyuridine (BrdU; Sigma B5002; 0.8 mg/ml) was provided in the drinking water for a period of 2 weeks, starting either 14 d before or 3 d after RMS AAV injection.

### Sensory manipulation

Naris plugs were made using a polytetrafluoroethylene (PTFE) hollow tube (inner diameter (ID) 0.3 mm; outer diameter (OD) 0.6 mm; VWR International) filled with suture thread knotted three times around shredded filaments of dental floss (Cummings *et al*., 1997). The mice were briefly anaesthetised using an isoflurane chamber and the Vaseline-lubricated naris plug (∼5-7 mm length) was completely inserted into the right nostril until the nostril eventually closed back behind the plug. For sham controls, the mice were also anaesthetised, and the plug was inserted into the right nostril and then immediately removed. After either procedure, mice were culled 24 hours later.

### Transcardial perfusions and immunohistochemistry

For tissue fixation, mice were anaesthetised with an intraperitoneal pentobarbital overdose. Breathing, paw-pinch and tail and ocular reflexes were all absent before starting transcardial perfusions. The mice were secured in place and the heart was exposed. A needle connected to a peristaltic pump was inserted into the left ventricle and the right atrium was cut to open the circulation. The following solutions were transcardially perfused: 25 ml of phosphate-buffered saline (PBS) with heparin (1:1000) was used to flush out the blood from the circulation, followed by 25 ml of 4 % paraformaldehyde (PFA; in 3 % sucrose, 60 mM PIPES, 25 mM HEPES, 5 mM EGTA and 1 mM MgCl_2_). The OBs were dissected and kept in 4 % PFA with PIPES for 1 d, then washed three times in PBS (five minutes each wash) and kept at 4 °C in PBS with 0.02 % sodium azide. The bulbs were embedded in 5 % agarose and cut coronally into 50 µm slices using a vibratome (VT1000S Leica).

Free floating slices were kept at 4 °C in PBS with 0.02 % sodium azide. For blocking and permeabilization of the slices, 10 % normal goat serum (NGS) was made in TritonX/PBS/azide (0.25 % TritonX; 0.02 % sodium azide). The slices were incubated in this solution for 2 h at room temperature floating on a shaker. Then, the slices were incubated with the primary antibodies (Table 1) overnight at 4 °C. The following day, the primary antibodies were removed, and the slices were washed three times with PBS, before being incubated with appropriate secondary antibodies (AlexaFluor) diluted in 10 % NGS, for 2 hours at room temperature. Labelled slices were mounted on glass slides with FluorSave mounting medium and covered with an appropriate glass coverslip. To permit BrdU detection, slices were first incubated in 2 M HCl for 30 min at 37 °C, washed thoroughly in PBS and then processed for immunohistochemistry as described above.

**Table 1.** Primary antibodies used.

| Antigen | Species | Dilution | Supplier and identifier |
| --- | --- | --- | --- |
| BrdU | Rat | 1 : 200 | Serotec OBT0030<br>RRID: AB_609568 |
| Calbindin | Rabbit | 1 : 5000 | Swant CB38a<br>RRID: AB_10000340 |
| Calretinin | Mouse | 1 : 5000 | Millipore MAB1568<br>RRID: AB_94259 |
| GAD67 | Mouse | 1 : 500 | Millipore MAB351R<br>RRID: AB_94905 |
| GFP | Chicken | 1 : 1000 | Abcam ab13970<br>RRID: AB_300798 |
| tdTomato | Rabbit | 1 : 1000 | Clontech 632496<br>RRID: AB_10013483 |
| TH | Mouse | 1 : 500 | Millipore MAB318<br>RRID: AB_2313764 |

### Imaging

OB fixed slices were imaged using a laser scanning confocal microscope (Zeiss LSM 710). Images were acquired with appropriate excitation and emission filters, a 1 AU pinhole aperture and a x40 oil immersion objective. For assessment of co-label, laser power and gain were adjusted manually to facilitate visualization of cells with low immunofluorescence signals. GFP co-label with other markers was assessed using Zen (Zeiss) and/or ImageJ (Fiji) software. Ambiguous co-label was assessed using orthogonal projections from z-stacks.

### Acute Slice Electrophysiology

DAT-tdTomato mice were used for electrophysiological recordings at either 5 or 9 weeks-post-injection (wpi). The mice were deeply anaesthetised with isoflurane and decapitated. Their OB was quickly extracted in ice-cold sucrose slicing medium (in mM: 240 sucrose, 5 KCl, 1.25 Na_2_HPO_4_, 2 MgSO_4_, 1 CaCl_2_, 26 NaHCO_3_ and 10 D-Glucose, bubbled with 95% O_2_ and 5 % CO_2_; unless otherwise noted, all reagents were from Sigma) and was sliced into 300 µm horizontal sections using a Leica vibratome (VT1000S). The sections were then placed in standard artificial cerebrospinal fluid (aCSF) (in mM: 124 NaCl, 2.5 KCl, 1.25 NaH_2_PO_4_, 2 MgSO_4_, 2 CaCl_2_, 26 NaHCO_3_ and 15 D-glucose, bubbled with 95 % O_2_ and 5 % CO_2_), for 30 min at 32-34 °C and at least 30 min at room temperature before being used for recordings.

The slices were held in place using a harp (Warner Instruments) and were recorded in oxygenated aCSF continuously perfused at a rate of ∼2 ml/min, warmed to physiologically relevant temperature (34 ± 2 °C) using an in-line heater (TC-344B, Warner Instruments). Ionotropic synaptic receptor blockers (10 µM NBQX, 50 µM APV, 10 µM SR-95531 (gabazine)) were added to the perfusion bath to block glutamatergic and GABAergic synaptic inputs.

Live-imaged dopaminergic cells were visualised using an upright microscope (Axioscop Eclipse FN1, Nikon) with a 40x water immersion objective and captured with a DMx 31BF03 camera (Scientifica). The fluorescence of tdTomato-positive and/or GFP-positive cells was revealed by CooLEDpe-100 light sources coupled with appropriate excitation and emission filters. Whilst resident dopaminergic neurons were identified by their expression of tdTomato and not GFP, adult-born dopaminergic cells were identified by their GFP expression, with or without co-expression of tdTomato. GFP fluorescence was much brighter than that of tdTomato and therefore, unlike tdTomato, could be visualised deeper in the slice. For this reason, targeting of adult-born dopaminergic neurons for electrophysiological recordings relied exclusively on their GFP expression. The identity of either tdTomato-positive or GFP-positive cells was confirmed by the appearance of fluorescence in the patch pipette and by its disappearance from the cell itself after membrane rupture. Data from resident tdTomato-positive cells were combined into a single group comprising recordings at both 5 and 9 wpi.

For whole cell recordings, borosilicate glass pipettes (OD 1.5 mm, ID 0.84 mm; WPI) were pulled using a vertical P10 puller (Narishige) with a resistance of 3.5-7.5 MΩ. Pipettes were filled with intracellular solution containing, in mM: 124 K-gluconate, 9 KCl, 10 KOH, 4 NaCl, 10 HEPES, 28.5 Sucrose, 4 Na_2_ATP, 0.4 Na_3_GTP (pH with KOH). We did not correct for a calculated liquid junction potential of ∼13 mV (https://swharden.com/software/LJPcalc/app/).

Patch-clamp recordings were performed using a MultiClamp 700B amplifier (Molecular Devices) and digitised using an Axon Digidata 1550B (Molecular Devices). Data were collected using Clampex software and analysed using customised MatLab scripts. Fast capacitance was compensated in the on-cell configuration after reaching a GΩ seal. Current clamp recordings were corrected for both bridge balance and pipette capacitance neutralization (set at 5.5 pF). Passive membrane properties were determined from a test pulse consisting of a 10 ms hyperpolarising voltage step (Δ-10 mV) from a holding voltage (V_hold_) of -60 mV. The recordings were sampled at 50 kHz, and Bessel-filtered at 10 kHz. The test pulse was performed before and after each protocol. Series resistance (R_s_), calculated from the peak current, was assessed continuously and recordings were excluded if they exceeded 30 MΩ at any point or changed more 20 % during any recording. No significant differences in R_s_ were observed in the recordings between any experimental group.

### Passive membrane properties

Resting membrane potential was recorded in current clamp (*I* = 0) soon after membrane rupture (<30 s) to minimise the effects of dialysing the intracellular solution, and was calculated as the mean voltage measured over 5 s. The resting membrane potential of spontaneously spiking cells was excluded from our analyses.

Input resistance (R_i_) and membrane capacitance (C_m_) were measured from the test pulse in voltage clamp. R_i_ was calculated from the difference in steady-state current evoked by a 10 ms hyperpolarizing step. C_m_ was calculated as the area under the transiently decaying curve from the peak of the current at the beginning of the voltage step to the steady-state current.

### Single action potential properties

Single action potentials were evoked by injecting 10 ms current steps of increasing amplitude starting from +0 pA in increments of at least Δ+10 pA from -60 mV. Current steps were injected until an action potential (V_m_>0 mV) was fired within the current step duration. From this protocol multiple characteristics of the action potential waveform were calculated: maximum rate of action potential rise (Max dV/dt) was measured as the largest positive rate of change (dV/dt) measured during the action potential; voltage threshold for firing an action potential (V_thresh_) was measured as the voltage at which the rate of action potential rise exceeded 10 V/s; maximum amplitude of the action potential (V_max_) was measured as the maximum voltage of the action potential; action potential width at half height (whh), was calculated at the midpoint between V_thresh_ and V_max_. Recordings were sampled at 200 kHz and smoothed using a 20 point (100 µs) sliding filter prior to differentiation for dV/dt.

Other single action potential features (current rheobase, latency to first spike, fast afterhyperpolarization) were measured from the first spike elicited by injecting increasing current steps of at least Δ+5 pA starting from 0 pA from -60 ± 3 mV for 500 ms. Current rheobase was calculated as the minimum current value at which the first action potential was fired; latency to the first spike was calculated as the time between the start of the current injection and the maximum voltage of the first spike; fast hyperpolarization (fAHP) was measured as the difference between spike voltage threshold and the minimum voltage after the first spike.

### Multiple spiking

Multiple spiking was elicited by 500 ms current injections, which were stopped when the cell reached depolarisation block. These recordings were used to determine both the single action potential features above (current rheobase, latency to first spike, fAHP) and multiple spiking properties (input-output relationship, instantaneous frequency (IF), variability in spike timing (coefficient of variation; CV), spike frequency adaptation (SFA), spike amplitude adaptation (SAA) and spike width adaptation (SWA)).

For the input-output relationship, the number of action potentials fired was calculated for each current step. Input-output functions were then described using various single-value parameters designed to simplify comparisons between experimental groups.

First, inter-spike interval (ISI) was calculated as the time between the V_max_ of two consecutive spikes in the 500 ms trace. ISIs were measured between all the spikes in a given trace and at each current level. This measure was used for calculating IF, CV and SFA. These parameters were measured for each sweep and calculated from the following formulas:

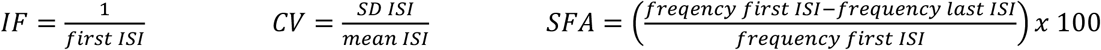

The V_max_ and whh of the first and last spike of the 500 ms trace were also measured for each sweep. These measures were used to calculate SAA and SWA from the following formulas:

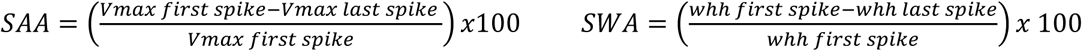

For IF and SAA, input-output curves were generated by plotting IF or SAA at each current level. Then, they were measured by applying a linear fit to the input-output plots to 0-80 % of the maximum spike number. For the other multiple spiking parameters such as CV, SFA and SWA, the relationship between these parameters and the injected current was not clearly linear. Thus, a single representative value for these measures was chosen at 1.5x rheobase in order to normalise for variability in initial cell responses to current injections.

### Medium afterhyperpolarization (mAHP)

In order to record the mAHP after a train of spikes, 10 action potentials were elicited by injecting 2 ms suprathreshold current injections (500 pA) at 50 Hz and the protocol was repeated over 10 sweeps. The amplitude of the mAHP was measured from the mean trace, and was calculated as the difference between V_hold_ and the minimum voltage in the 250 ms following the end of the spike train. Any trace that did not successfully produce 10 action potentials in response to the 10 stimulation pulses was not included in the average trace for analysis.

### Sag potential

Sag potential was measured from 500 ms hyperpolarising current steps (at least -Δ10 pA) from V_hold_ until the steady state voltage (V_ss_), corresponding to the end of the current step, reached at least – 100 mV. The sag index was then calculated with the following formula:

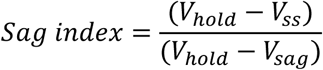

where V_ss_ represents the steady state voltage, and V_sag_ represents the minimum voltage during the current step. In order to calculate the sag index at ∼-100 mV, a linear interpolation method was used (Chittajallu *et al*., 2013).

### Quantitative PCR (qPCR)

All experimental steps for qPCR were performed in compliance with the MIQE guidelines (Bustin *et al*., 2009) and as previously described (Byrne *et al*., 2022). Adult (>2-month-old) wild-type mice were sacrificed by cervical dislocation followed by decapitation. The brain was quickly removed and placed in cold PBS. The right OB was dissected and placed in RNAlater (Thermo Fisher Scientific) overnight at 4 °C and stored at – 20 °C until further processing. Total RNA extraction was performed using the RNAqueous® Micro Kit (Life Technologies). RNA was eluted in 20 μl of elusion buffer and quantified using Nanodrop (Thermo Fisher Scientific) and Qubit (RNA HS Assay Kit, Life Technologies). A total of 300–500 ng of total RNA were used for cDNA synthesis with SuperScript™ II Reverse Transcriptase (Life Technologies), 0.5 mM dNTPs (Life Technologies), 5 μM OligodT23VN (IDT), 10 μM DTT (Life Technologies) and 0.2 Units RNase inhibitor (SUPERase•In™, Invitrogen). RNA was denatured for 5 min at 65 °C, before adding the reaction mix. Retrotranscription was performed at 42 °C for 90 min, followed by 20 min at 70 °C for enzyme inactivation. Standard RT-PCRs were run for all samples, including non-template controls, to corroborate amplification of single bands of the appropriate size. Cycling conditions were 95 °C for 3 min, (95 °C, 15 s; 65–55 °C [-0.5 °C per cycle], 20 s; 68 °C, 25 s), 20 cycles, (95 °C, 15 s; 55 °C, 20 s; 68 °C, 25 s), 20 cycles, 68 °C, 5 min, 4 °C, hold. Reactions were run on a SimpliAmp™ Thermal Cycler (Applied Biosystems). Primers used were ActB_For, cctctatgccaacacagtgc; ActB_Rev, cttctgcatcctgtcagcaa; Th_For, tgcctcctcacctatgcact; Th_Rev, gtcagccaacatgggtacg. Primers were designed using the Universal Probe Library Assay Design Centre (Roche), compatible with UPLs 157 (ActB) and 3 (Th).

qPCR reactions were run using the FastStart Essential DNA Probes Master mix (Roche) on the LightCycler ® 96 Real-Time PCR System (Roche). Final primer concentration was 0.5 μM, and probe concentration was 0.1 μM. Cycling conditions were 95 °C for 10 min, (95 °C, 10 s; 60 °C, 30 s), 55 cycles, 4 °C, hold. Technical duplicates were performed for all samples and reactions. ActB was used as housekeeping reference gene for Th expression quantification. Amplification efficiency for all primer pairs was determined by obtaining a standard curve. Briefly, the cDNAs from each cDNA synthesis reaction were pooled and serially diluted. qPCR reactions were performed as above. Ct amplification curve values were plotted as a function of dilution factor and the amplification efficiency determined from the slope of the linear regression as in E = 10^(-1/slope)^-1. Amplification efficiency (E ± SE) for ActB primers was 1.07 ± 0.02, for Th primers, 1.01 ± 0.03. Fold changes in Th expression levels were obtained using the 2^(–ddCt)^ method (Livak & Schmittgen, 2001; Taylor *et al*., 2019), first normalising Ct levels for each sample to housekeeping Ct levels and subsequently normalising to the average of the control (sham) conditions.

### Statistical analysis

Statistical analysis was carried out using Prism (Graphpad), Matlab (Matworks) or SPSS (IBM). Data are reported as mean ± SEM; n and N refer to the number of cells and mice, respectively. All tests were two-tailed, with α set at 0.05. Details of all individual statistical tests are reported alongside the appropriate results. Data were assessed for normality using the D’Agostino and Pearson omnibus test where datasets were sufficiently large, or the Shapiro-Wilk test if not. Parametric or non-parametric tests were used for normally distributed and non-normally distributed data, respectively. For dual-factor analyses, data were analyzed with 2-way ANOVAs. Since there is no available non-parametric alternative to 2-way ANOVA, data that were not normally distributed by D’Agostino and Pearson omnibus test or Shapiro-Wilk test were analyzed for normality using a custom-written Matlab script which fitted a normal distribution and calculated the percentage of dataset variance accounted for by that fit. If >10 % of the variance could not be explained by the fit, the data were transformed by standard methods (detailed in each case in the Results) and re-tested for normality. Then, a 2-way ANOVA was used to analyze data rendered normal by the above methods. When no common transform resulted in normally distributed data, a 2-way ANOVA was performed on data ranks (Akritas, 1990). After 1-way ANOVA, post-hoc pairwise comparisons were performed using Tukey’s or Dunn’s multiple comparisons tests for parametric and non-parametric tests, respectively. After 2-way ANOVA, post-hoc pairwise comparisons were performed using Sidak’s multiple comparisons test, with the p values adjusted for three comparisons (5 wpi sham vs 5 wpi occluded; 9 wpi sham vs 9 wpi occluded; resident sham vs resident occluded). Fisher’s exact test was performed when analysing proportions; where multiple pairwise Fisher’s exact tests were carried out on the same data, α was Bonferroni-corrected as indicated in the text.

## Results

### Selective labelling of adult-born OB dopaminergic neurons

Investigating the functional maturation and plasticity of adult-born OB dopaminergic neurons required a means of specifically labelling those cells to enable visually targeted patch-clamp recordings in the acute slice preparation. We achieved this specificity through a combinatorial approach: using viral vector injections into the adult RMS to specifically target recently born neuroblasts migrating towards the OB, and doing so in a conditional transgenic line that restricted virally encoded fluorophore expression to cells with dopaminergic identity.

After injecting AAV-CAG-FLEx-GFP into the RMS of > 2-month-old DAT*^IREScre^* x Rosa26-floxed-stop-tdTomato (DAT-tdTomato) mice and waiting 5 or 9 wpi (Figure 1A) we observed GFP-expressing cells in the OB whose spatial distribution and morphology was entirely consistent with an adult-born origin and dopaminergic identity. In fixed OB tissue, GFP-positive cells were scattered rather evenly and sparsely throughout the glomerular layer and outer external plexiform layer (Figure 1B). This is consistent with initial infection of a subset of migrating neuroblasts that later take up residency throughout the OB, and reflects the expected location of bulbar dopaminergic neurons (Hökfelt *et al*., 1975; Banerjee *et al*., 2015; Kosaka *et al*., 2019; Byrne *et al*., 2022). GFP-positive cells also had morphological features consistent with dopaminergic identity, including a small soma and dendrites that ramified locally within the glomerular neuropil (Figure 1C).

**Figure 1.**
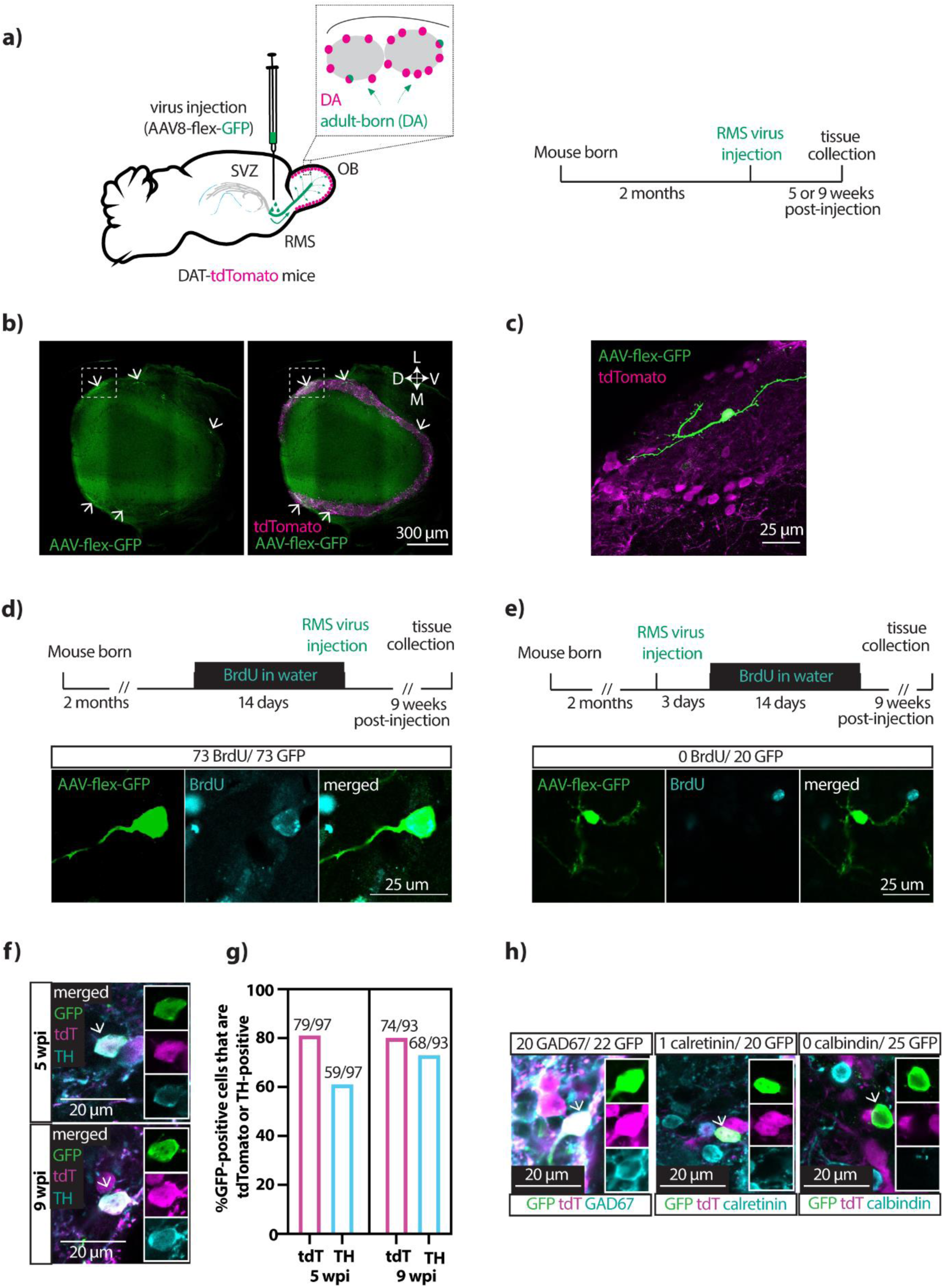
Labelling specificity for adult-born dopaminergic neurons. **a)** Diagram showing the viral injection strategy and the experimental timeline used to label adult-born dopaminergic neurons in the OB. The inset shows magnification of two glomeruli representing the types of labelled cells. SVZ, subventricular zone; RMS, rostral migratory stream; DA, dopaminergic. **b)** Single-plane stitched image showing sparse label of GFP+ cells (white arrows) distributed around the glomerular layer of the OB. Scale bar, 300 µm. L, lateral; M, medial; D, dorsal; V, ventral. **c)** Magnification of the inset area shown with dashed lines in **b)**, showing single-cell morphology of a GFP-expressing glomerular layer neuron. **d)** Timeline of RMS virus injections *after* 2-week administration of BrdU in drinking water. Below, single plane images show an adult-born GFP+ neuron co-labelled with BrdU. Scalebar, 25 µm. **e)** Timeline of RMS virus injections *before* 2-week administration of BrdU in drinking water. Below, single plane images show an adult-born GFP+ neuron which does not co-label with BrdU. Scale bar, 25 µm. **f)** Single-plane confocal images showing GFP+ cells in the glomerular layer of the OB at 5 wpi and 9 wpi. Insets show single-channel fluorescence. Both cells, indicated with arrows, are also positive for tdTomato and TH. Scalebar, 20 µm. **g)** Quantification of the percentages of GFP+ cells in the glomerular layer that expressed tdTomato or TH at 5 wpi and 9 wpi. **h)** Single-plane confocal images of GFP-positive neurons in the glomerular layer of the OB (white arrows), in tissue co-labelled for GAD67, calretinin or calbindin. Insets show single-channel fluorescence. Left, a GFP-, tdT- and GAD67-positive cell; middle, a GFP- and tdT-positive but calretinin-negative cell; right, a GFP-positive cell that is negative for both tdT and calbindin. Scalebar, 20 µm.

We confirmed that our GFP-positive cells were adult-born by administering the thymidine analogue BrdU via drinking water for the 2 weeks immediately prior to AAV injection (Figure 1D). Consistent with their production from cells that divided during recent BrdU availability, we found that every GFP-expressing cell in OB tissue fixed at 9 wpi was BrdU-positive (Figure 1D; n = 73/73 GFP-positive cells; N = 3 mice). To place a lower bound on GFP-positive cell age, we also administered BrdU via drinking water for 2 weeks starting 3 d *after* AAV injection (Figure 1E). At 9 wpi, we found no glomerular layer cells that were double-labelled for both BrdU and GFP (Figure 1F; n = 0/20 GFP-positive cells; N = 3 mice). This demonstrates that GFP-expressing OB neurons were never the result of mitotic divisions occurring after AAV injection. Moreover, the combined results from both BrdU experiments show that the cohort of GFP-positive neurons present in the OB at any given timepoint was entirely generated in a temporally restricted manner over, at the very most, a ∼2 week period prior to AAV injection.

Specificity for dopaminergic identity was assessed by labelling GFP-positive neurons for cell-type-specific markers. We observed high levels of tdTomato co-label, as expected given the reliance of both fluorophores on DAT-Cre expression (Figure 1F,G; 5 wpi: n = 79/97 = 81 %, N = 7 mice; 9 wpi: n = 74/93 = 80 %, N = 5 mice; Fisher’s exact test, p = 0.85). The dopamine-synthesising enzyme TH is a specific marker for dopaminergic neurons in the OB (Hökfelt *et al*., 1975; Rosser *et al*., 1986), and we saw a slight, non-significant increase between 5 wpi and 9 wpi in the proportion of GFP-positive cells that were also TH-positive (Figure 1F,G; 5 wpi: n = 59/97 cells = 61 %; N = 7 mice; 9 wpi: n = 68/93 cells = 73 %; N = 6 mice; Fisher’s exact test, p = 0.09). By 9 wpi, the percentage of TH-positive GFP-expressing neurons was very near the range of TH co-label previously reported for OB cells in different DAT-Cre lines (75-85 %; Banerjee *et al*., 2015; Vaaga *et al*., 2017; Byrne *et al*., 2022). Consistent with OB dopaminergic cells’ ability to co-release GABA, we also found that GFP-positive cells co-labelled for the GABA-synthesising enzyme GAD67 (Figure 1H; n = 20/22 cells = 91 %, N = 3 mice; Kiyokage *et al*., 2010; Kosaka *et al*., 2019). In contrast, we saw almost no co-expression in GFP-positive cells of the two markers for other, non-dopaminergic GABAergic neurons in the bulb’s glomerular layer: either calbindin (Figure 1H; n = 0/25 cells = 0 %, N = 3 mice) or calretinin (Figure 1H; n = 1/20 cells = 4 %, N = 3 mice; Kohwi *et al*., 2007; Kosaka & Kosaka, 2007; Panzanelli *et al*., 2007; Parrish-Aungst *et al*., 2007).

### Functional identity of live-labelled adult-born OB dopaminergic neurons

Our approach produced live-labelled GFP-positive cells that could be visualised in acute OB slices and targeted for patch-clamp recordings (Figure 2A,B). Using endogenous GFP fluorescence in live tissue, we were able to detect the occasional GFP-positive cell in the OB’s glomerular layer from 2 wpi. However, these cells remained very dimly labelled and extremely rare between 2-4 wpi, preventing reliably productive patch recordings from being obtained at these timepoints. From 5 wpi, we observed brightly labelled GFP-positive cells at higher densities and were able to record from them on a regular basis. In parallel, we also obtained recordings from GFP-negative, tdTomato-positive resident dopaminergic neurons.

**Figure 2.**
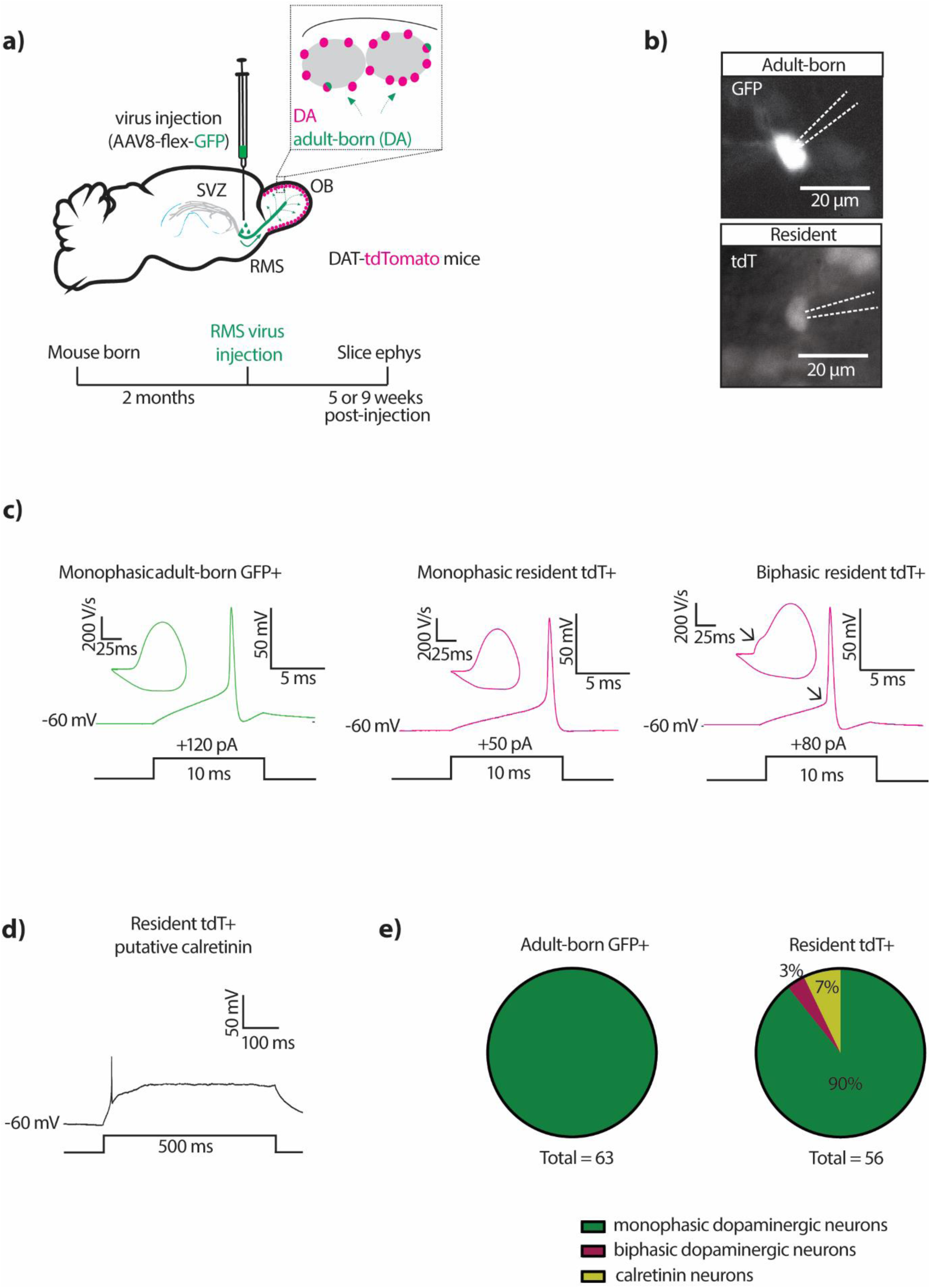
Functional identity of live-labelled adult-born OB dopaminergic neurons **a)** Diagram showing the viral injection strategy and the experimental timeline used to live-label adult-born dopaminergic neurons in acute OB slices. Ephys, electrophysiology; all other conventions as in Figure 1A. **b)** Example epifluorescence images of endogenous GFP and tdTomato expression in a patched adult-born and resident neuron, respectively. Dashed lines show pipette location; scalebar, 20 µm. **c)** Example traces of single action potentials and their respective phase plane plots fired at current threshold to 10 ms duration somatic current injection, in a 5 wpi adult-born GFP-positive neuron (green trace, left) and two resident tdTomato (tdT)-positive neurons (magenta traces, right). Arrows point to the sharp ‘kink’ at spike onset and its corresponding ‘bump’ in the rising phase of the phase plane plot of the biphasic resident action potential. **d)** Example trace of a characteristic single spike and subsequent plateau potential in response to 500 ms current injection in a resident tdTomato-positive putative calretinin-expressing cell. **e)** Pie charts showing the proportions of recorded adult-born GFP-positive cells (left) and resident tdTomato-expressing cells (right) that displayed monophasic dopaminergic, biphasic dopaminergic or calretinin-like electrophysiological properties.

The functional features of our recorded GFP-positive cells were consistent with their being adult-generated and dopaminergic. All dopaminergic OB neurons produced in adulthood are anaxonic, forming part of a morphologically defined subtype which is associated with a characteristic ‘monophasic’ action potential waveform (Galliano *et al*., 2018, 2021; Dorrego-Rivas *et al*., 2024; Lau *et al*., 2024). Indeed, all GFP-positive neurons we recorded had monophasic spikes (Figure 2C,E). This was not due to an inability to find or identify axon-bearing dopaminergic ‘biphasic’ action potentials in our adult slice preparation, because such waveforms were occasionally clearly observed in recordings from resident neurons (Figure 1C,E; cells with biphasic spikes were removed from our dataset). In addition, GFP-expressing cells lacked other functional features associated with non-dopaminergic identity. Some DAT-tdTomato cells are known to co-express calretinin and have the characteristic ‘immature-like’ functional features of calretinin-expressing glomerular interneurons (Fogli Iseppe *et al*., 2016; Benito *et al*., 2018; Byrne *et al*., 2022). We did see a small proportion of resident tdTomato-positive neurons with these properties (these cells were also removed from our dataset), but no GFP-positive cells behaved this way (Figure 2D,E). Overall, these data, coupled with the fixed tissue analyses presented above (Figure 1) show that our live labelling approach attained a high level of specificity for adult-generated OB dopaminergic neurons.

### Passive membrane properties of adult-born OB dopaminergic neurons are largely mature by 5 weeks post-injection

Having established the adult-born and dopaminergic specificity of AAV- and DAT-Cre-driven GFP expression, we went on to obtain a set of whole-cell patch-clamp recordings from GFP-positive neurons in acute OB slices taken at either 5 or 9 wpi. These groups were chosen based on the earliest timepoint at which reliable data could be gathered from GFP-labelled OB neurons (5 wpi), as well as previous findings indicating significant adult-born glomerular layer cell maturation and plasticity over a similar 1-2 month period of cell age (∼5-9 wpi; Winner *et al*., 2002; Kohwi *et al*., 2007; Livneh *et al*., 2014; Kovalchuk *et al*., 2015; Maslyukov *et al*., 2018). Recordings from GFP-negative, DAT-tdT-positive resident neurons were taken as readouts of established, presumed mature functionality (Banerjee *et al*., 2015; Galliano *et al*., 2018; Lau *et al*., 2024).

Initial assessment of passive membrane properties found that GFP-positive neurons at 5 wpi were indistinguishable from those recorded at 9 wpi, and that both adult-generated groups were almost identical to resident dopaminergic neurons. Input resistance and membrane capacitance did not differ across the three groups (Figure 3A-C; Table 2). Resting membrane potential, on the other hand, was more hyperpolarised in adult-born dopaminergic neurons compared to resident cells (Figure 3D; Table 2).

**Figure 3:**
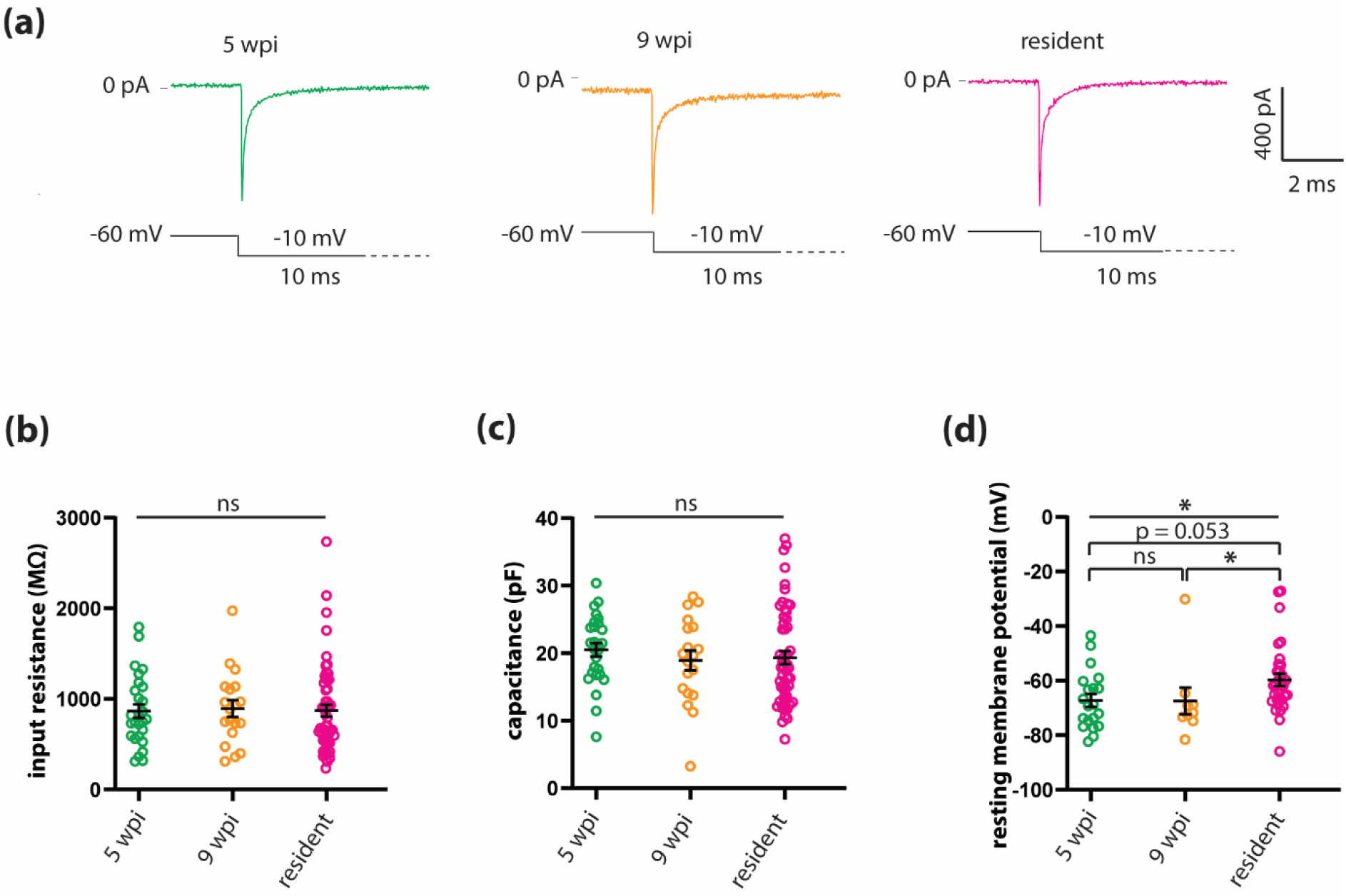
Passive membrane properties of adult-born OB dopaminergic neurons are largely mature by 5 weeks post-injection. **a)** Example traces of voltage clamp membrane test series from 5 wpi (green), 9 wpi (orange), and resident (magenta) dopaminergic neurons. **b, c, d)** Quantification of input resistance (**b**), membrane capacitance (**c**) and resting membrane potential (**d**), recorded from 5 wpi (green), 9 wpi (orange) and resident (magenta) dopaminergic neurons. Lines show mean ± SEM; each dot shows data from one cell; straight lines report overall result of one-way ANOVA or non-parametric equivalent; brackets show results of post-hoc pairwise tests; *, p < 0.05.

**Table 2.**
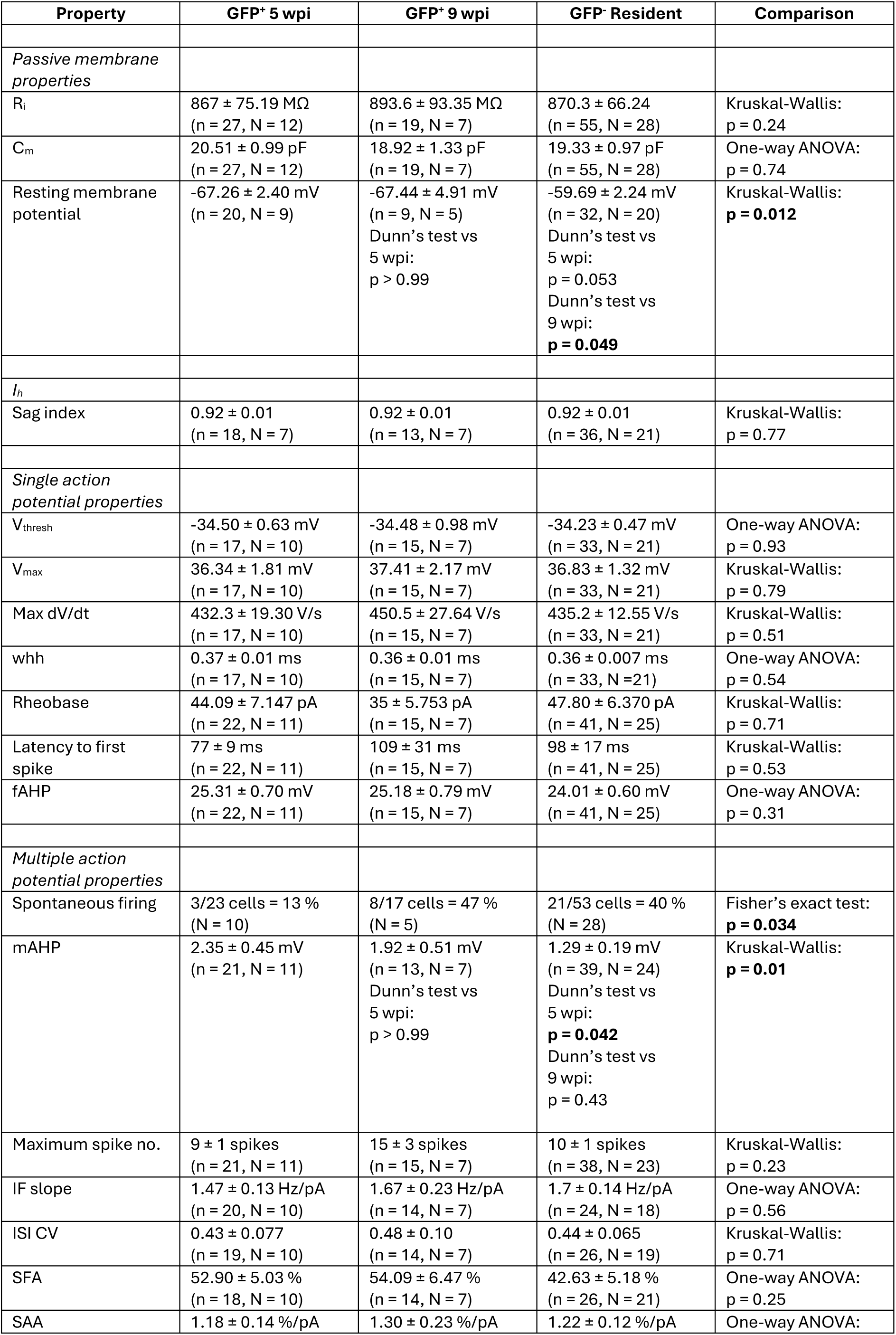
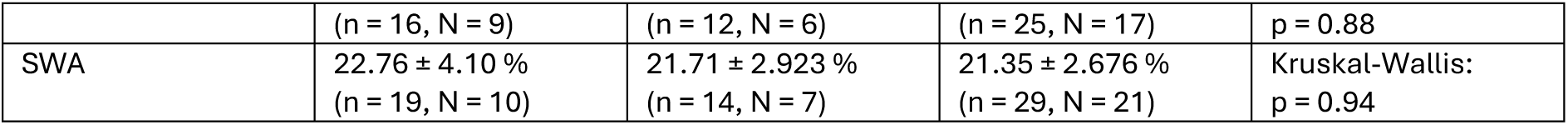
Maturation of electrophysiological properties in adult-born OB dopaminergic neurons. All descriptive data and statistical comparisons involve cells from sham-treated mice only. Values show mean ± SEM n, number of cells; N, number of mice; IF, instantaneous frequency; ISI, interspike interval; CV, coefficient of variation; SFA, spike frequency adaptation; SAA, spike amplitude adaptation; SWA, spike width adaptation. Bold values show significant effects.

### Single action potential properties are fully mature in adult-born OB dopaminergic cells by 5 weeks post-injection

We first explored active intrinsic physiological properties by recording and analysing single action potentials fired at current threshold in response to a brief 10 ms current injection step (Figure 4A). In all respects, we found that single spike features were completely indistinguishable between adult-born neurons recorded at 5 wpi and 9 wpi and resident dopaminergic cells (Table 2). In particular, at the earliest possible timepoint we saw fully mature action potentials in terms of voltage threshold (Figure 4B), maximum voltage (Figure 4C), maximum rate-of-rise (Figure 4D), and spike width (Figure 4E).

**Figure 4.**
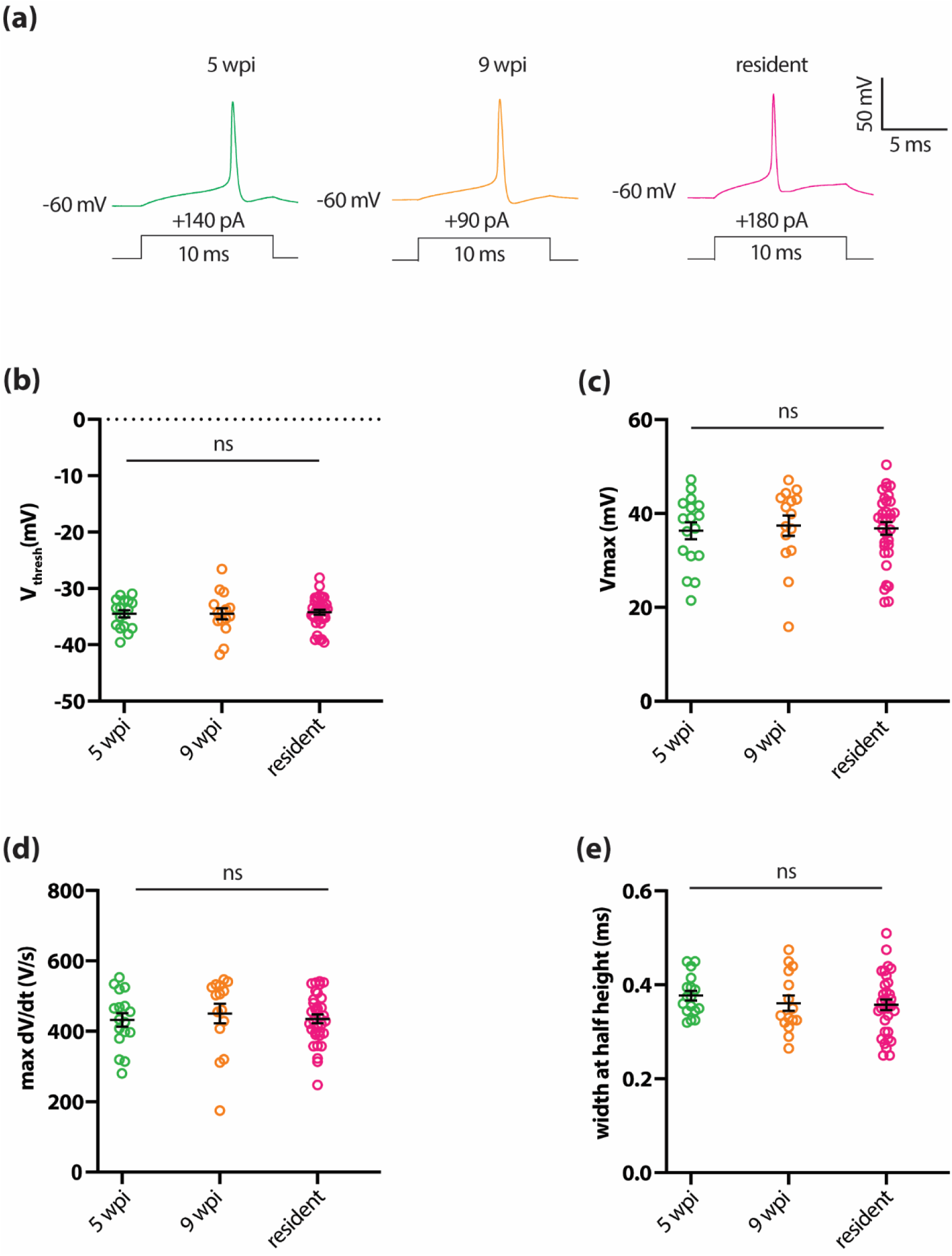
Single action potential properties are fully mature in adult-born OB dopaminergic cells by 5 weeks post-injection**. a)** Example traces of single action potentials elicited in current clamp by 10 ms threshold somatic current injection in 5 wpi (green), 9 wpi (orange) and resident (magenta) dopaminergic neurons. **b-e)** Quantification of voltage threshold (**b**), maximum voltage (**c**), maximum rate of rise (**d**), width at half height (**e**). All conventions as in Figure 3.

### Spontaneous firing and medium afterhyperpolarisation undergo significant and independent maturation in adult-born OB dopaminergic neurons

Despite the fully mature nature of individual action potentials in 5 wpi GFP-positive cells, we found that the probability of spontaneous firing varied significantly with maturation (Table 2), with more immature adult-born neurons being less likely to spike under *I* = 0 current clamp conditions (Figure 5A,B). Given that these recordings were obtained in the presence of blockers for all major ionotropic neurotransmitter receptors in OB circuits (see Methods), this immature functional feature in 5 wpi cells was most likely due to their having distinct intrinsic properties. Differences in resting membrane potential could potentially underlie the maturational increase in spontaneous spiking, but we could not directly test this because we were unable to reliably record resting membrane potential in spontaneously spiking cells. Nevertheless, in the data we were able to obtain from non-spontaneously active cells (Figure 3D), resting membrane potential was not significantly higher in more mature adult-born neurons.

**Figure 5.**
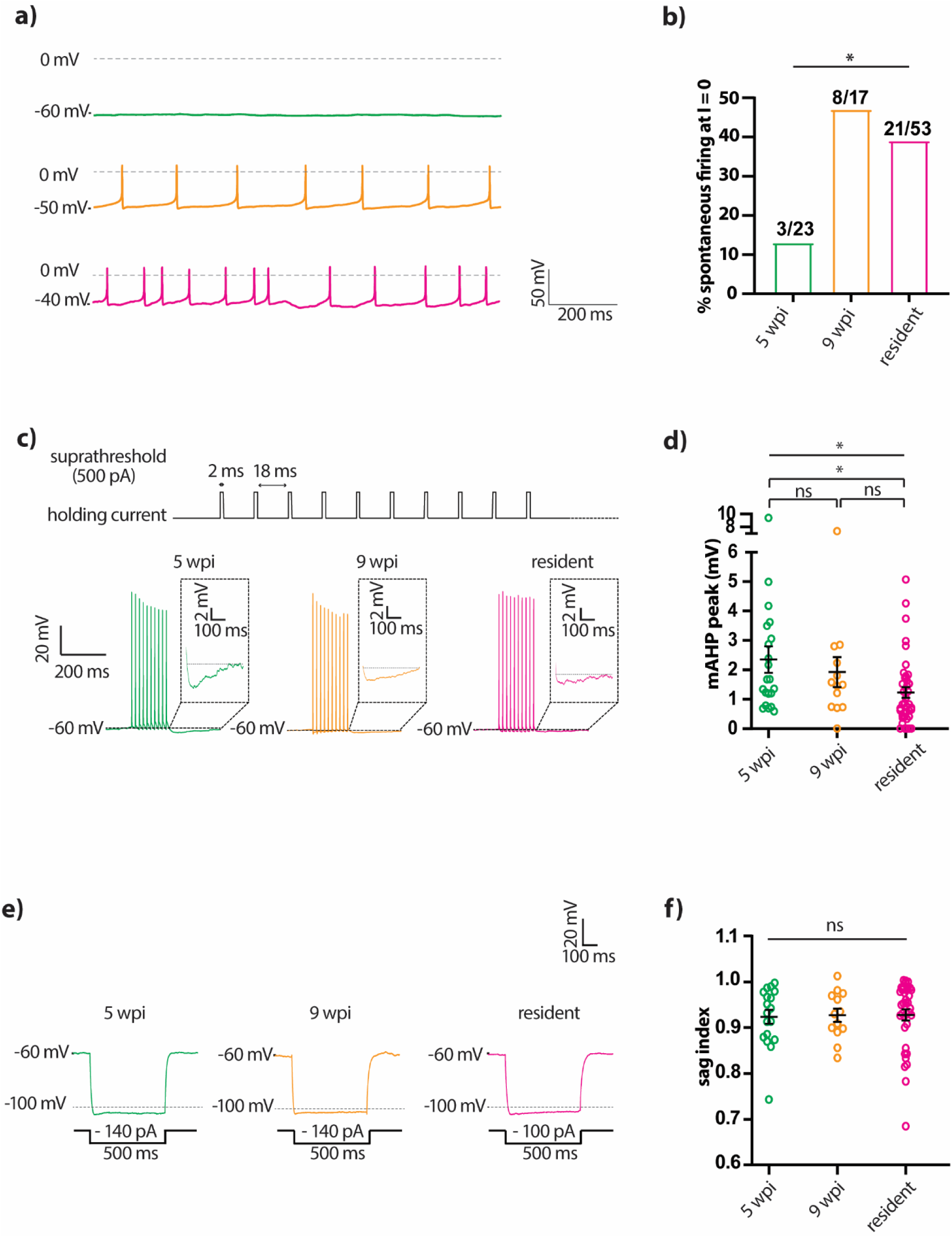
Spontaneous firing and medium afterhyperpolarisation undergo significant maturation in adult-born OB dopaminergic neurons**. a)** Example individual traces of spontaneous firing recorded in the absence of current injection (*I* = 0) from 5 wpi (green), 9 wpi (orange), and resident (magenta) dopaminergic neurons. **b)** Bar plot shows the percentage of spontaneously active neurons at *I* = 0 in each group. Fisher’s exact test: *, p < 0.05. **c**) Top, current clamp protocol for medium afterhyperpolarization (mAHP) consisting of 2 ms suprathreshold current injections at 50 Hz. Bottom, example traces of the mAHP in 5 wpi, 9 wpi and resident dopaminergic neurons. Straight dashed lines indicate the baseline voltage for each trace. Insets show mAHP magnification. **d)** Quantification of peak mAHP amplitude. All conventions as in Figure 3. **e)** Example traces of sag potential elicited by hyperpolarizing current steps in 5 wpi, 9 wpi and resident dopaminergic neurons. **f)** Quantification of sag index. All conventions as in Figure 3.

One additional intrinsic property that differed significantly across 5 wpi, 9 wpi and resident groups was the amplitude of the medium afterhyperpolarisation (mAHP), which we measured immediately following a brief train of ten spikes evoked at 50 Hz (Figure 5C). Despite considerable variation within groups, mAHP amplitude decreased with cell maturation, and was significantly greater in 5 wpi adult-born, GFP-positive dopaminergic neurons compared to resident tdTomato-positive cells (Figure 5D; Table 2). However, mAHP amplitude did not differ significantly between spontaneously active and inactive neurons (2-way ANOVA on square root plus one transformed data; effect of maturation F_2,61_ = 7.53, p = 0.0012; effect of spontaneous firing F_1,61_ = 0.053, p = 0.82; interaction F_2,61_ = 2.57, p = 0.085), so these two maturational changes are unlikely to be closely interrelated. Similarly, we found no significant within-group correlations between mAHP amplitude and resting membrane potential (5 wpi: Spearman r = - 0.17, p = 0.55, n = 15 cells, N = 9 mice; 9 wpi: Spearman r = 0.086, p = 0.92, n = 6 cells, N = 5 mice; resident: Spearman r = -0.33, p = 0.14, n = 21 cells, N = 20 mice).

In many cell types, both spontaneous firing and the mAHP, as well as the resting membrane potential, can be regulated by the *I*_h_ current carried by hyperpolarisation-activated cyclic nucleotide-gated (HCN) cation channels (Maccaferri *et al*., 1993; Maccaferri & McBain, 1996; Oswald *et al*., 2009). OB dopaminergic cells can possess an *I*_h_ current which influences resting membrane potential, though its small amplitude means that it does not directly contribute to spontaneous pacemaking activity in these neurons (Pignatelli *et al*., 2013). We measured the sag index as a proxy for *I*_h_ amplitude and observed small but detectable sag potentials in most OB dopaminergic neurons, as previously described (Figure 5E,F; Pignatelli *et al*., 2013; Lau *et al*., 2024). We found that sag index was fully mature in 5 wpi GFP-positive neurons, with no difference in this measure between groups (Figure 5E,F; Table 2). We also found no significant within-group correlations between sag index and mAHP amplitude (5 wpi: Spearman r = -0.19, p = 0.51, n = 14 cells, N = 12 mice; 9 wpi: Spearman r = -0.6, p = 0.073, n = 10 cells, N = 7 mice; resident: Spearman r = -0.25, p = 0.23, n = 25 cells, N = 28 mice), nor between sag index and resting membrane potential (5 wpi: Spearman r = -0.40, p = 0.14, n = 15 cells, N = 9 mice; 9 wpi: Spearman r = -0.54, p = 0.30, n = 6 cells, N = 5 mice; resident: Spearman r = -0.13, p = 0.54, n = 24 cells, N = 20 mice). Sag index also did not significantly differ between spontaneously spiking and silent cells (2-way ANOVA; effect of maturation F_2,57_ = 0.20, p = 0.82; effect of spontaneous firing F_1,57_ = 0.045, p = 0.83; interaction F_2,57_ = 0.35, p = 0.71). Overall, this strongly suggests that *I*_h_ variation did not contribute to any of the maturational effects we observed in other intrinsic properties.

### Multiple spiking properties are fully mature in adult-born OB dopaminergic cells by 5 weeks post-injection

The late maturation of the mAHP in adult-born dopaminergic neurons (Figure 5C,D) might be predicted to regulate different features of multiple spike firing (Duménieu *et al*., 2015; Dwivedi & Bhalla, 2021). We therefore performed a comprehensive analysis of the spike rates and patterns elicited by prolonged 500 ms current injection steps in adult-born and resident OB dopaminergic neurons. However, we found no differences between groups in any measure of evoked multiple spike firing (Figure 6; Table 2). In particular, maximum spike number (Figure 6B) and average input-output functions (Figure 6C; mixed model 2-way ANOVA, effect of maturation: F2, 192 = 1.10, p = 0.33) were fully mature by 5 wpi.

**Figure 6.**
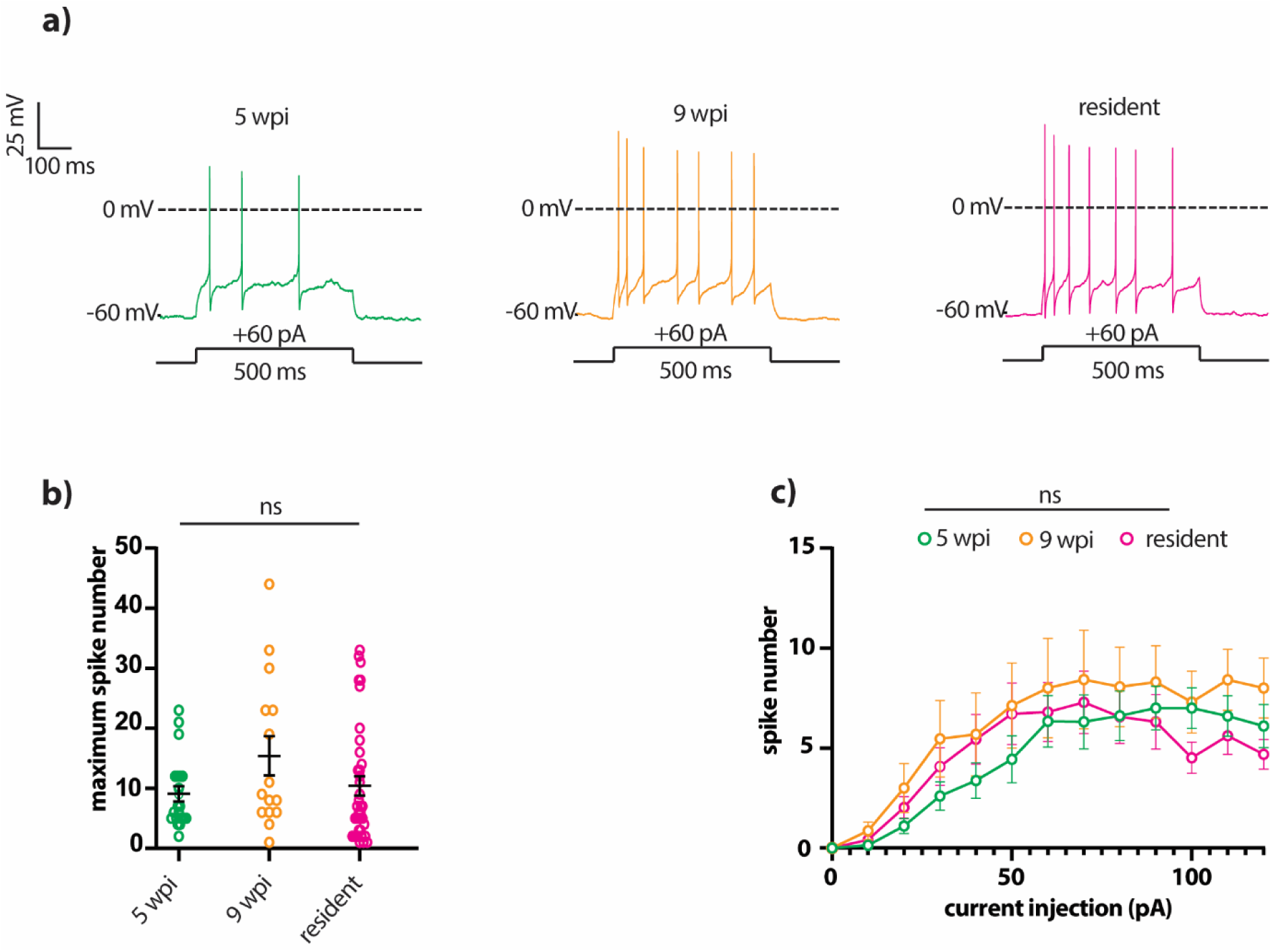
Evoked multiple spiking properties are fully mature in adult-born OB dopaminergic neurons by 5 wpi. **a)** Example single traces of suprathreshold multiple firing elicited by 500 ms somatic current injections (+60 pA) into 5 wpi (green), 9 wpi (orange), and resident (magenta) dopaminergic neurons. **b)** Quantification of the maximum number of action potentials fired to any input stimulus. All conventions as in Figure 3. **c)** Mean ± SEM action potential number fired to each input stimulus. Mixed-model 2-way ANOVA, effect of maturation: ns, non-significant.

### Brief sensory deprivation induces spike amplitude increases in all OB dopaminergic neurons

Immature adult-born neurons have been shown to have an elevated capacity for activity-dependent plasticity in many morphological and functional features (Schmidt-Hieber *et al*., 2004; Ge *et al*., 2007; Tashiro *et al*., 2007; Kelsch *et al*., 2009; Nissant *et al*., 2009; Gu *et al*., 2012; Livneh *et al*., 2014; Alvarez *et al*., 2016). We tested whether this was also true of intrinsic electrophysiological properties in adult-generated OB dopaminergic neurons, using a physiologically relevant, brief period of sensory deprivation via 24 h unilateral naris occlusion (Figure 7A). We previously employed this manipulation to successfully induce different forms of plasticity in OB dopaminergic cells in juvenile mice (Galliano *et al*., 2021; Byrne *et al*., 2022), and here confirmed its efficacy in adult animals by replicating the downregulation produced by 24 h occlusion in whole-bulb *Th* mRNA expression (Byrne *et al*., 2022; Figure 7B; sham, mean ± SEM = 1.03 ± 0.12, N = 5; occluded, 0.69 ± 0.052, N = 5; t-test, t_8_ = 2.57, p = 0.033).

**Figure 7.**
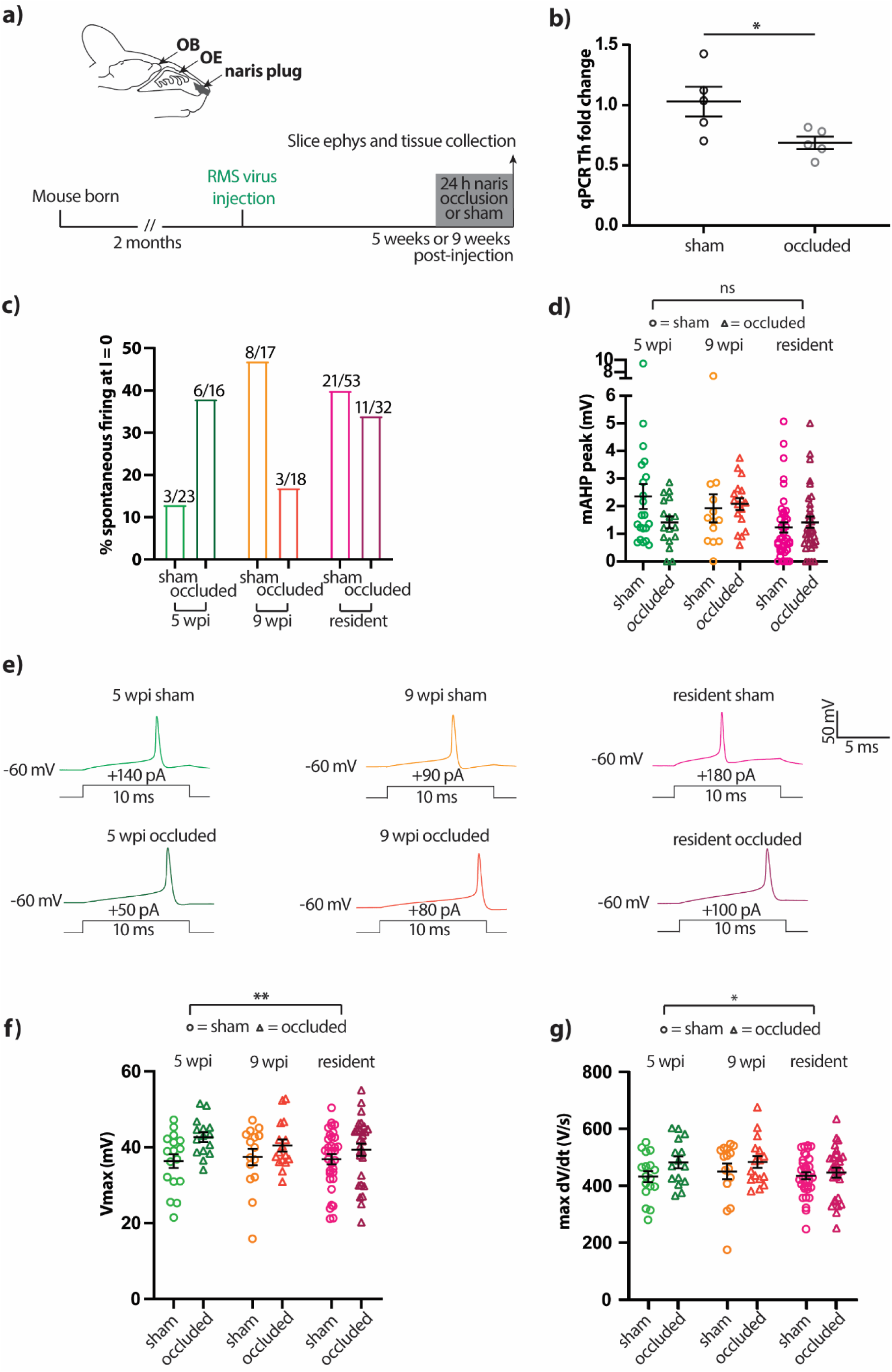
Brief sensory deprivation induces spike amplitude increases in all OB dopaminergic neurons. **a)** Diagram and timeline of unilateral naris occlusion for 24 h sensory deprivation. OE, olfactory epithelium; ephys, electrophysiology. **b)** Mean ± SEM fold change vs control for whole-OB *Th* qPCR. Each dot shows data from one mouse; t-test: *, p < 0.05. **c)** Percentage of neurons with spontaneous firing across all timepoints and treatment groups. **d)** mAHP amplitude across all timepoints and treatment groups. Each symbol shows data from one cell, lines show mean ± SEM. Effect of occlusion in 2-way ANOVA: ns, non-significant. **e)** Example traces of single action potentials elicited in current clamp by 10 ms threshold somatic current injection in 5 wpi (green), 9 wpi (orange) and resident (magenta) dopaminergic neurons in sham (light) or occluded (dark) treatment groups. **f,g)** Quantification of maximum voltage (**f**) and maximum rate of rise (**g**). All conventions as in **d)**; *, p < 0.05; **, p < 0.01.

Although we observed differences between 5 wpi, 9 wpi and resident control groups in both spontaneous spiking (Figure 5B) and mAHP amplitude (Figure 5D), these properties were unaffected by 24 h sensory deprivation (Figure 7C,D; Table 3). Indeed, across the full range of intrinsic functional features we examined in adult-born and resident OB dopaminergic cells, almost none were impacted in any way by brief naris occlusion, and we never saw any evidence for significantly elevated plasticity in adult-born cells in general, or younger 5 wpi adult-born cells in particular (Table 3).

**Table 3.**
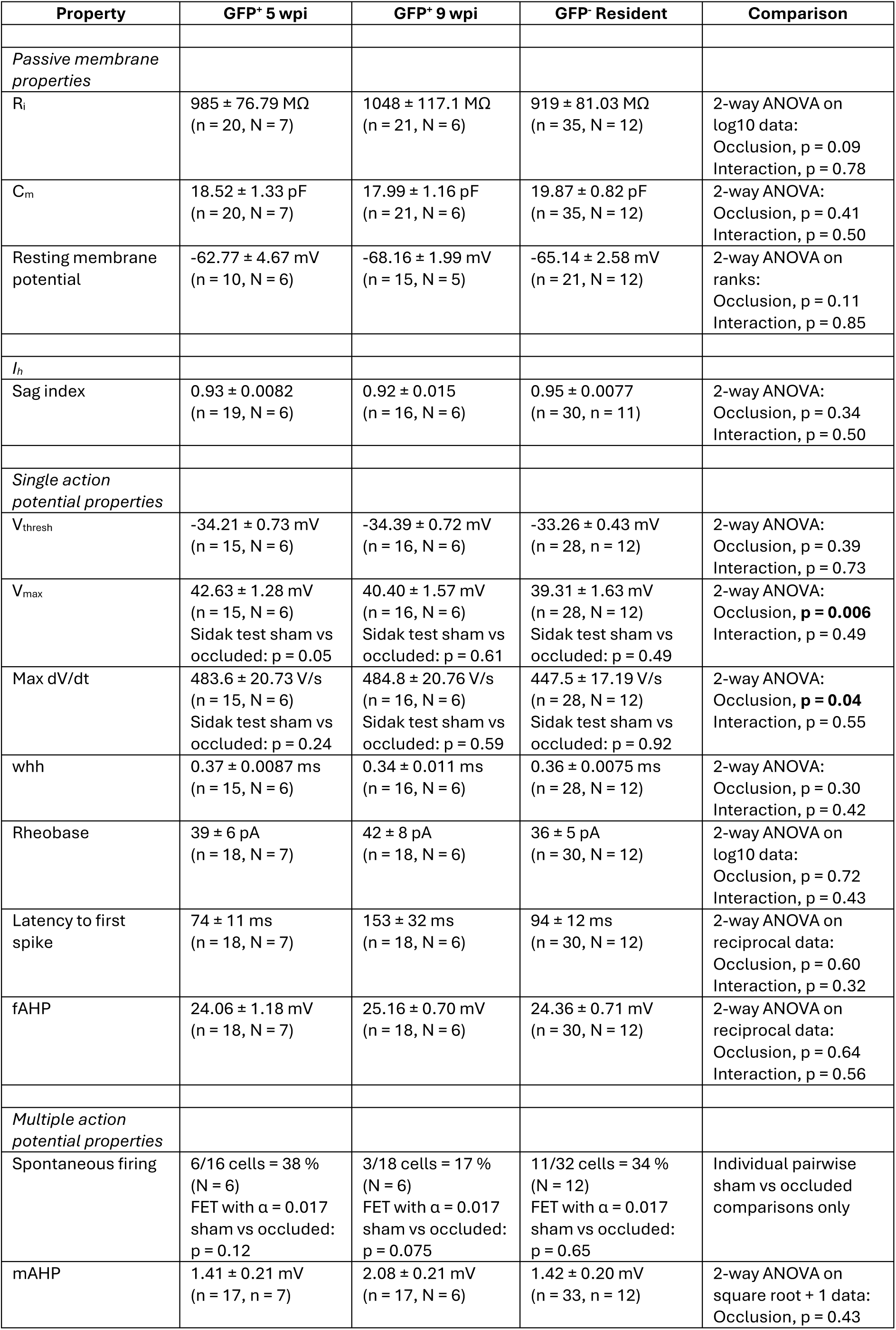
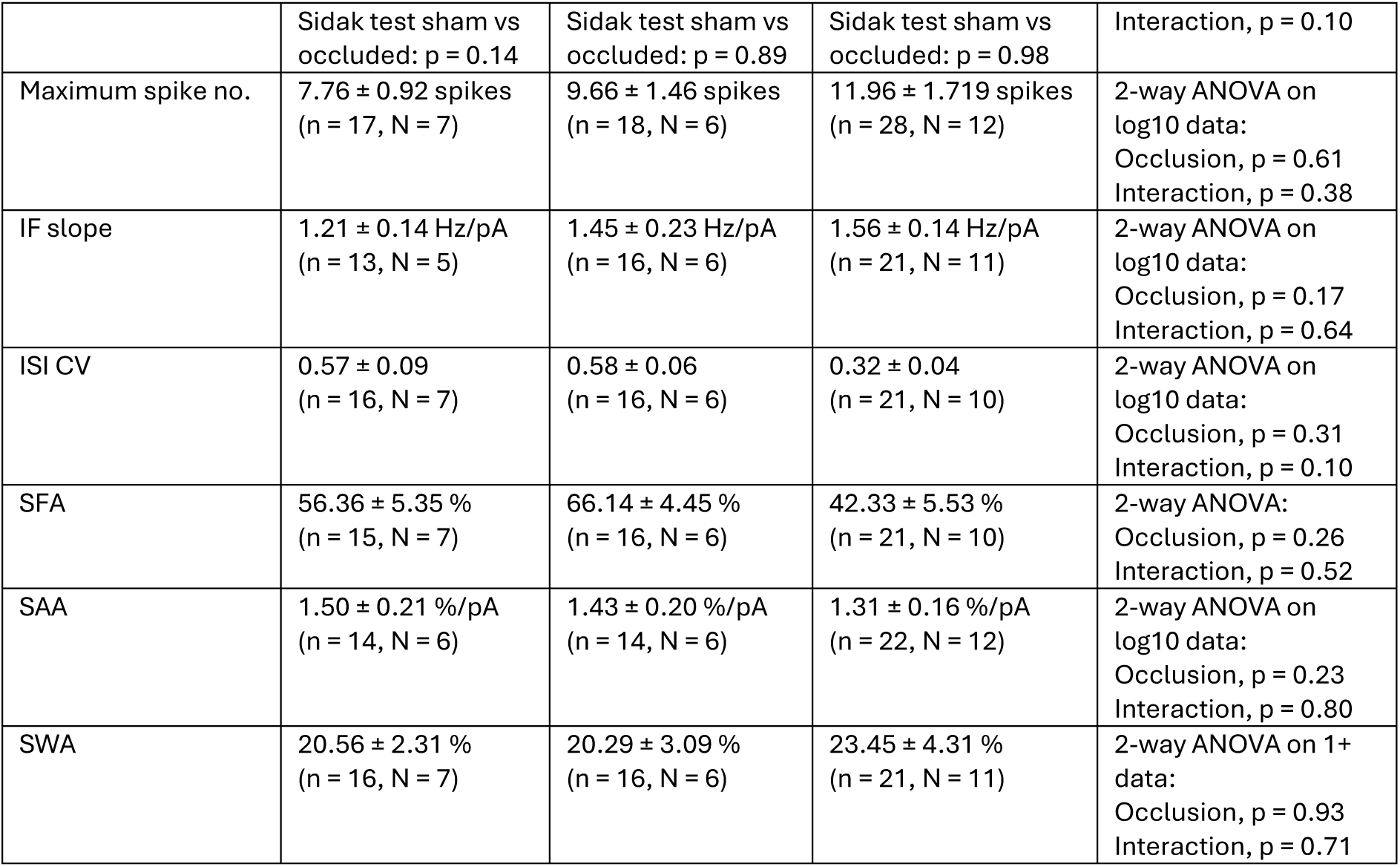
Effect of brief sensory deprivation on electrophysiological properties of adult-born OB dopaminergic neurons. All descriptive data shown are from 24 h unilateral naris-occluded mice; statistical comparisons also include sham data from Table 2. Values show mean ± SEM; ‘Interaction’ refers to the effect of occlusion x maturation; n, number of cells; N, number of mice; IF, instantaneous frequency; ISI, interspike interval; CV, coefficient of variation; SFA, spike frequency adaptation; SAA, spike amplitude adaptation; SWA, spike width adaptation; FET, Fisher’s exact test. Bold values show significant effects.

We did, however, uncover significant activity-dependent plasticity in two closely related features of the action potential waveform. After 24 h unilateral naris occlusion, spike maximum voltage and maximum rate-of-rise were both significantly increased across all groups – adult-born and resident neurons alike (Figure 7E-G; Table 3). These effects tended to be stronger in adult-generated dopaminergic cells, and especially in the more immature 5 wpi GFP-positive group (Figure 5F,G; Table 3), but we found no statistically significant age-x-occlusion interactions (Table 3). Functional plasticity of intrinsic electrophysiological properties therefore exists in this specific subtype of adult-born OB neuron, but is not restricted to cells at any particular maturational stage.

## Discussion

Here we used a conditional targeting strategy to successfully and selectively label a specific, dopaminergic subclass of adult-born OB periglomerular neurons for electrophysiological characterisation at different stages of their maturation. Live GFP label driven by DAT-Cre expression first became reliably visible around 5 weeks of cell age, and by this stage GFP-positive adult-born neurons had almost entirely mature intrinsic electrophysiological properties. We saw evidence for maturation only in the incidence of spontaneous firing, which became more prevalent in 9-week-old neurons, and in mAHP amplitude, which decreased with cell maturity. Brief sensory deprivation via 24 h unilateral naris occlusion revealed no sign of elevated plastic potential in immature or mature adult-born dopaminergic cells. However, measures of action potential amplitude and rate-of-rise were consistently increased after occlusion, regardless of neuronal maturation state.

### Selective live label for a specific subtype of adult-born OB periglomerular cells

Our conditional live label approach successfully achieved selectivity on two separate levels. First, temporal specificity: targeting migrating neuroblasts with a single AAV injection into the RMS ensured virally mediated expression only in adult-generated OB cells within a restricted cellular age range. Second, cell-type specificity: Cre-dependent GFP expression in these adult-born neurons was restricted to dopaminergic cells expressing Cre under the control of the DAT promoter. GFP expression was therefore highly selective to adult-born dopaminergic cells of a restricted age cohort, though a small degree of off-target labelling is always unavoidable in such conditional expression approaches (Fischer *et al*., 2019). In general, our strategy mirrored similar targeting of specific subtypes and cohorts of adult-born OB granule cells, an approach which has revealed distinct functional features in these neuronal subclasses (Malvaut *et al*., 2017; Hardy *et al*., 2018). Importantly, we note that while AAV infection itself has been reported to be detrimental to the health and survival of some adult-born cell populations (Johnston *et al*., 2021) the almost identical functional resemblance we found between AAV-labelled adult-born cells and their resident neighbours shows that AAV-infected adult-generated OB dopaminergic neurons are capable of entirely healthy, normal maturation.

### Rapid maturation of intrinsic functional properties in adult-born OB dopaminergic neurons

Achieving cell-type specificity via DAT-Cre expression had the drawback that adult-born dopaminergic neurons could only be reliably identified from around 5 weeks post-AAV injection. As soon as we were able to consistently record from GFP-positive cells at this timepoint, their intrinsic functional properties were almost entirely mature. This is consistent with the early establishment of functional properties reported for postnatally born OB periglomerular cells in general, which fire action potentials, receive olfactory nerve inputs, and display almost fully mature odorant-evoked calcium responses just 1-2 weeks after they arrive in the glomerular layer (Belluzzi *et al*., 2003; Grubb *et al*., 2008; Kovalchuk *et al*., 2015). Furthermore, while major morphological changes occur over time in adult-born periglomerular neurons, no significant alterations in dendritic morphology have been found within the ∼1-2-month time period we studied here (Livneh *et al*., 2014; Su *et al*., 2023). Later maturation has been found in the sensory response properties of adult-born periglomerular cells in general, which spike more strongly and respond to a broader range of odorant stimuli at 4 weeks compared to 8 weeks after RMS injection (Livneh *et al*., 2014). Our findings, however, suggest that such later refinements are either largely driven by changes in non-dopaminergic periglomerular neurons, and/or mainly depend upon maturational alterations in connectivity (e.g. Livneh *et al*., 2009) rather than intrinsic properties. Similar explanations must also pertain to the distinct odorant response properties found for mature adult-born versus resident periglomerular cells (Fomin-Thunemann *et al*., 2020) – given our findings here, these differences are unlikely to be driven by the intrinsic electrophysiological properties of the dopaminergic periglomerular subtype.

The rapid establishment of intrinsic physiology in adult-born OB dopaminergic neurons contrasts with their relatively delayed maturation of full dopamine signalling capacity. Although dopaminergic fate is determined early via gene expression patterns established in regions of the subventricular zone (Hack *et al*., 2005; Merkle *et al*., 2007; de Chevigny *et al*., 2012; Coré *et al*., 2020), and although Th mRNA is expressed while these cells are still migrating to the glomerular layer (Baker *et al*., 2001; Pignatelli *et al*., 2009), expression of TH protein does not reach mature levels until several weeks later (Winner *et al*., 2002; Kohwi *et al*., 2007; Kovalchuk *et al*., 2015). Our findings suggest that DAT-driven Cre expression does not begin in most adult-born dopaminergic neurons until they have been in the OB’s glomerular layer for several weeks, and that expression of TH protein in Cre-positive neurons takes even longer (Figure 1). The adult-born OB ‘dopaminergic’ cell type is therefore functionally mature in many respects, and likely capable of participating in odour-driven information processing, long before it has the necessary machinery to release dopamine itself. The function of such a potentially extended period of neurochemical maturation is unclear, but may be related to the renowned activity-dependence of dopamine-synthesising enzymes in this cell type (Nadi *et al*., 1981; Baker *et al*., 1983; Bonzano *et al*., 2016; Galliano *et al*., 2021; Byrne *et al*., 2022). Perhaps only the most well-integrated, strongly activated adult born neurons over an extended timeframe are able to reach full dopaminergic functional capacity.

### Later maturation of select intrinsic properties

Between 5-week-old, 9-week-old and resident OB dopaminergic neurons, we did find some evidence for intrinsic physiological maturation. The incidence of spontaneous spiking increased with maturation over this later time period, an observation generally in line with the increased spontaneous firing frequency observed for generally labelled adult-born periglomerular cells between ∼1-2 weeks and 8 weeks of age (Kovalchuk *et al*., 2015), although in vivo sodium channel-dependent spontaneous calcium transients in such cells appear to decrease in frequency between ∼3-4 weeks and 6 weeks of maturation (Maslyukov *et al*., 2018). Spontaneous pacemaking activity is a characteristic functional feature of OB dopaminergic neurons, known to be controlled by persistent sodium and T-type voltage-gated calcium currents (Pignatelli *et al*., 2005; Pignatelli & Belluzzi, 2017). It will be crucial in future work to determine how the maturation of these or other contributing currents allow spontaneous firing to become prevalent in adult-born dopaminergic cells.

We also found that the amplitude of the mAHP decreased during later maturation of adult-generated dopaminergic neurons. This is consistent with a similar age-dependent decrease observed in developing OB mitral cells, which was found to be mainly driven by alterations in apamin-sensitive calcium-dependent potassium currents (Duménieu *et al*., 2015). OB dopaminergic neurons do possess some calcium-dependent potassium currents (Pignatelli *et al*., 2005), which may therefore be a prime candidate underlying mAHP changes, either via direct changes in the maturational expression of the channels themselves (Cingolani *et al*., 2002) or via indirect alterations in the calcium entry that activates them. The mAHP can also be influenced by *I*_h_, which is present in this cell type (Pignatelli *et al*., 2013). However, we found no evidence for maturation of the *I*_h_-dependent sag potential, making it unlikely that *I*_h_ changes could be producing the smaller mAHP seen in older adult-born neurons. The functional implications of this maturational mAHP decrease also remain unclear – changes in the mAHP can strongly influence spike firing patterns (Duménieu *et al*., 2015; Dwivedi & Bhalla, 2021), but here we found no accompanying maturational alterations in multiple spiking (Figure 6). Perhaps, although significant, the mAHP decrease over time in adult-born OB dopaminergic cells was not sufficiently large to impact on spiking behaviour. Alternatively, the influence of lower mAHP in more mature neurons might require more naturally patterned input stimuli to make its impact on spike patterning apparent.

### No elevated plastic potential in immature adult-born OB dopaminergic cells

Immaturity is often associated with increased plasticity in adult-generated neurons (Schmidt-Hieber *et al*., 2004; Ge *et al*., 2007; Tashiro *et al*., 2007; Kelsch *et al*., 2009; Nissant *et al*., 2009; Gu *et al*., 2012; Livneh *et al*., 2014; Alvarez *et al*., 2016), but here we found no evidence for specific or heightened forms of intrinsic plasticity in immature OB dopaminergic cells. This may be due to the duration of the manipulation we employed – although brief 24 h sensory deprivation was sufficient to decrease whole-OB Th mRNA in adult animals (Figure 7B) and was able to produce cell subtype-specific plastic alterations in OB dopaminergic cells in juvenile mice (Galliano *et al*., 2021; Byrne *et al*., 2022), longer periods of occlusion may be needed to induce specific forms of plasticity in adult-born neurons. However, in studies of generally labelled adult-generated periglomerular cells, even weeks of naris occlusion had no effect on migration, morphology, or spontaneous or odour-evoked calcium responses (Mizrahi, 2007; Li *et al*., 2023). Perhaps other forms of activity manipulation may have revealed greater plastic changes? Indeed, in generally labelled adult-born periglomerular neurons, cell-intrinsic silencing via early overexpression of potassium channels is associated with significantly decreased migration, survival, morphological complexity and odour-evoked responses (Li *et al*., 2023, but see Bugeon *et al*., 2021), while prolonged odour enrichment can produce increased synaptogenesis, altered dendritic arborisations, and distinct odour response profiles (Livneh *et al*., 2009, 2014; Livneh & Mizrahi, 2011). Applying such manipulations, or brief naris occlusion, earlier in the lifetime of an adult-born dopaminergic cell may also have revealed elevated plastic potential, though this was not possible in the current study due to the relatively late onset of DAT-dependent live label. Finally, we assessed only intrinsic properties in the present work, so it is still possible that brief sensory deprivation could produce marked plasticity in the synaptic inputs and/or outputs of immature adult-generated OB dopaminergic neurons.

### Intrinsic action potential plasticity in all OB dopaminergic neurons

Across all groups of OB dopaminergic cells, regardless of maturation state, we did find consistent plastic alterations in action potential height and speed induced by brief 24 h naris occlusion. Given that all of the neurons in the present study had monophasic spike waveforms indicative of the anaxonic OB dopaminergic subtype (Chand *et al*., 2015; Galliano *et al*., 2018, 2021; Dorrego-Rivas *et al*., 2024; Lau *et al*., 2024), this contrasts with the lack of action potential plasticity seen in this subtype after 24 h naris occlusion in juvenile mice (Galliano *et al*., 2021). Animal age – juvenile vs adult – may be a factor here, but sample size is a more likely contributor. Compared to 124 cells here, only 24 monophasic DAT-tdT cells were recorded in the juvenile study, where trends were nevertheless seen towards larger maximum voltage and rate-of-rise after 24 h occlusion (Galliano *et al*., 2021).

What might be the mechanism underlying the altered spike waveform in adult OB dopaminergic neurons? Changes in the rising phase of the action potential are usually associated with voltage-gated sodium channel contributions around spike onset. Given that monophasic-firing cells lack an axon, and have none of the specialised spike initiation machinery usually found at the axon initial segment (Galliano *et al*., 2018), alterations in sodium channel spatial distributions or subcellular localisations are unlikely to drive the action potential plasticity we observed. Instead, in spikes that initiate somatically, it may be alterations in local sodium channel density and/or regulation that produce greater sodium influx once spikes have been triggered (Rama *et al*., 2015; Routh *et al*., 2017; Zbili *et al*., 2020).

Finally, the functional implications of faster and large action potentials for information processing in glomerular neurons remain entirely unclear, though fascinating to speculate upon. These changes could permit more reliable action potential propagation throughout the dendritic tree, though this propagation has been shown to be already extremely reliable under baseline conditions (Bywalez *et al*., 2016). They might also be associated with elevated calcium influx and more reliable vesicle exocytosis at presynaptic release sites (Zbili *et al*., 2020). In cells that provide local, intraglomerular dendritic release of both dopamine and GABA (Borisovska *et al*., 2013; Liu *et al*., 2016; Pignatelli & Belluzzi, 2017; Vaaga *et al*., 2017), either change might be expected to increase the contribution to local glomerular inhibition. On first inspection such increased inhibition may appear dysfunctional when OB networks are deprived of input. However, complex glomerular interactions, including between different types of GABA-releasing neurons (Toida, 2008; Banerjee *et al*., 2015; Parsa *et al*., 2015; Liu *et al*., 2016; Kiyokage *et al*., 2017; Liu, 2020) may mean that increased spike strength in the dopaminergic subtype could potentially result in overall decreased local levels of inhibition, and thereby contribute to compensatory changes that offset decreased external excitatory drive. Such considerations underscore even further the vital importance of taking into account cell subtype identity when aiming to understand the maturation, plasticity and functional roles of adult-born neurons.

## Acknowledgements

This work was supported by a H2020 European Research Council Consolidator Grant (725729; FUNCOPLAN), and BBSRC Research Grant (BB/N014650/1) to MSG. We wish to thank Elisa Galliano and Laura Andreae for comments on the manuscript, Juan Burrone and Sandrine Thuret for advice, and past and present members of the Grubb laboratory for technical assistance and stimulating discussion.

## Conflict of interest

The authors declare no conflicts of interest.

## Author contributions

CT, MC, ML, LPB and MSG designed the experiments; CT, MC and ML performed the experiments; CT, ML and MG analysed the data; CT and MG wrote the paper.

## Data availability statement

Upon publication, all primary data generated in this study will be made fully and openly available under a CC BY license (at https://doi.org/10.18742/27728289).

## Notes

### Competing Interest Statement

The authors have declared no competing interest.

